# Drought frequency predicts life history strategies in *Heliophila*

**DOI:** 10.1101/493270

**Authors:** J. Grey Monroe, Brian Gill, Kathryn G. Turner, John K. McKay

## Abstract

Explaining variation in life history strategies is a long-standing goal of evolutionary biology. For plants, annual and perennial life histories are thought to reflect adaptation to environments that differ in the frequency of stress events such as drought. Here we test this hypothesis in *Heliophila* (Brassicaceae), a diverse genus of flowering plants native to Africa, by integrating 34 years of satellite-based drought measurements with 2192 herbaria occurrence records. Consistent with predictions from classic life history theory, we find that perennial *Heliophila* species occur in environments where droughts are significantly less frequent compared to annuals. These associations are predictive while controlling for phylogeny, lending support to the hypothesis that drought related natural selection has influenced the distributions of these strategies. Additionally, the collection dates of annual and perennial species indicate that annuals escape drought prone seasons during the seed phase of their life cycle. Together, these findings provide empirical support for classic hypotheses about the drivers of life history strategy in plants - that perennials out compete annuals in environments with less frequent drought and that annuals are adapted to environments with more frequent drought by escaping drought prone seasons as seeds.

## Introduction

Understanding the causes and consequences of life history variation is a longstanding goal of ecology and evolutionary biology (Cole, 1954). In plants, life histories are especially diverse, with herbaceous species that complete their life cycle in a number of weeks to trees that live for thousands of years (Brown, 1996). Along this continuum in angiosperms an important division exists, distinguishing annuals which complete their seed to seed life cycle within a single calendar year from perennials which can persist over multiple years. Annual plants flower once, set seed, senesce, and then die, spending at least some portion of the year as a seed, where they are relatively protected from environmental stress. In contrast, perennial plants can continue vegetative growth after reproduction and must survive conditions experienced during all seasons. These represent fundamentally different life history strategies, but the ecological factors that explain their evolution and distributions remain empirically uresolved (Friedman & Rubin, 2015).

Classical theory predicts shorter life spans in environments where adult mortality is high (Charnov & Schaffer, 1973; Stearns, 1992; Franco & Silvertown, 1996). In plants, this has been extended to the hypothesis that annuality is adaptive when it allows plants to escape drought (Schaffer & Gadgil, 1975). Lack of water is perhaps the greatest threat to survival during vegetative or reproductive growth and annuals can remain dormant (and protected as a seed) during drought. Thus, environments with greater seasonal drought frequency may select for annual life histories that complete reproduction prior to drought prone seasons. Conversely, environments with less frequent drought may select for perennial species, which benefit from multiple bouts of reproduction and competitive advantage by preventing recruitment of annual species (Corbin & D’Antonio, 2004). These predictions have been supported by the observation of annuals in arid environments in *Oryza perennis* (Morishima *et al*., 1984) and *Oenothera* (Evans *et al*., 2005). Additionally, annual and perennial species of *Nemesia* were qualitatively associated with winter rather and summer rainfall environments respectively (Datson *et al*., 2008) and annual species of *Scorzoneroides* were associated with environments classified as unpredictable (Cruz-Mazo *et al*., 2009). However, whether the history frequency of drought events indeed predicts the distributions annual or perennial life history strategies has yet to be tested.

Here we combine a long-term global dataset of satellite detected drought events with metadata from natural history collections to test these classic hypotheses within the African endemic mustard genus, *Heliophila* L. (Brassicaceae). If annuality is an adaptive strategy allowing plants to escape drought prone seasons, then drought frequency should predict the distribution of life history strategies across landscapes, and annual species should be more commonly associated with drought prone regions than perennial species. Furthermore, if annual species have adapted to escape drought prone seasons, observations of growing annual species (i.e. occurring in forms other than seed) should be rare during drought prone seasons. Phylogenetic relatedness can influence tests of associations between species’ traits and their environments (Felsenstein, 1985; Barrett *et al*., 1996), and therefore we assessed the relationship between life history distribution and drought frequency in a phylogenetic context.

## Materials and Methods

### Data

#### Availability

All analyses were performed using R. All data and the source code to produce this manuscript are available at https://github.com/greymonroe/heliophila. Software used is listed in the supplement.

#### Satellite-detected drought data

Remotely sensed data is a powerful tool for characterizing seasonal patterns in drought because it is less limited in spatial and temporal scope and resolution than weather stations or field observations (AghaKouchak *et al*., 2015). To quantify the frequency of drought during different seasons across landscapes, we used the remotely sensed Vegetative Health Index (VHI), which measures landscape scale reductions in plant cover and temperature conditions characteristic of drought (Kogan, 2001). Generated from data collected by NOAA AVHRR satellites since 1981, the VHI combines Normalized Difference Vegetation Index (NDVI) derived measures of vegetative stress (Vegetative Condition Index - VCI) with temperature stress indicated by anomalies in thermal spectra (Temperature Condition Index - TCI). The VHI of year *y* during week *w* of [1, 52] at pixel *i* is derived from the following equations, where *n* is the number of years observed.

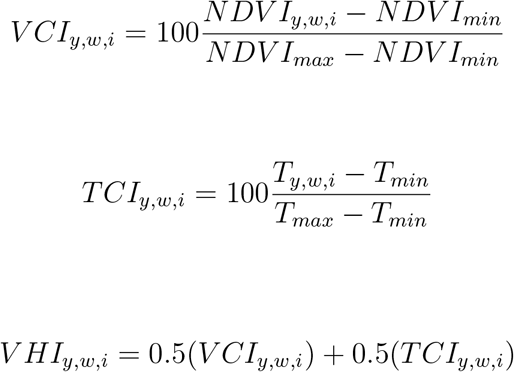

where *NDVI_min_* = *min*(*NDVI*_1981,*w,i*…_*NDVI*_1981+*n,w,i*_) and *NDVI_max_* = *max*(*NDVI*_1981,*w,i*…_*NDVI*_1981+*n,w,i*_) and *T_min_* = *min*(*T*_1981,*w,i*…_*T*_1981+*n,w,i*_) and *T_max_* = *max*(*T*_1981,*w,i*…_*T*_1981+*n,w,i*_)

Thus, VHI measurements are standardized according to conditions historically observed at each locations. These measurements have been validated and generally used for evaluating drought risk and predicting crop yields in agriculture (e.g., Rojas *et al*., 2011; Kogan *et al*., 2016). But they also present a new tool to study seasonal patterns in the frequency of drought across environments and to test hypotheses about the effect of drought on ecological and evolutionary processes (Kerr & Ostrovsky, 2003). As such, the VHI has been applied recently to study drought related ecology of natural species and proven useful for predicting intraspecific variation in drought tolerance traits and genes (Mojica *et al*., 2016; Dittberner *et al*., 2018; Monroe *et al*., 2018b). Here, we accessed VHI data at 16*km*^2^ resolution from 1981 to 2015 (https://www.star.nesdis.noaa.gov/smcd/emb/vci/VH/vh_ftp.php) to characterize the seasonal drought frequencies experienced by annual and perennial *Heliophila* species.

#### Life history data for *Heliophila*

*Heliophila* is a genus of flowering plants endemic to the southern portion of Africa including the Cape Floristic and Succulent Karoo Regions. These are among the most botanically diverse environments on Earth and the *Heliophila* species occurring there are considered to make up the most diverse genus of the family Brassicaceae (Mummenhoff *et al*., 2005; Mandáková *et al*., 2012). This genus includes both perennial and annual species and this change in life history strategy has likely arisen multiple independent times (Appel & Al-Shehbaz, 1997; Mummenhoff *et al*., 2005). Furthermore, the fine scale climatic heterogeneity of Southern Africa is ideal for studying the distribution of traits in relation to environmental parameters (Sayre *et al*., 2013). We used life histories reported by Mummenhoff *et al*. (2005), grouping species with annual or perennial life histories. Perenniality was defined based any form of perennial life history (e.g., herbs, shrubs, mixed, etc). Because the nature of species reported with mixed traits were unknown (i.e. plasticity vs. genetic variation), we classified these species here as perennial since they can maintain vegetative growth after reproduction at least to some capacity.

#### *Heliophila* occurrence records

Botanists have collected and maintained over 350 million botanical specimens worldwide over the past 300 years (Thiers, 2016). Herbarium specimens and their associated metadata have been used since the 1960s to study species’ geographical distributions (reviewed by Willis *et al*. (2017) and Lang *et al*. (2018)). And as they become digitized (Soltis, 2017), these collections have been used to study relationships between trait distributions, geography, and climate (Davis *et al*., 2015; Stropp *et al*., 2016; Wolf *et al*., 2016; Václavi’k *et al*., 2017). To characterize the distributions of annual and perennial *Heliophila* species, all records for the genus *Heliophila* were downloaded from the Global Biodiversity Information Facility (gbif.org) on July 21, 2018 (GBIF, 2018).

#### Sequence data for phylogeny

An alignment of ITS I and II sequences for *Heliophila* species was obtained from the authors of Mandáková *et al*. (2012). Individual ITS I and II sequences for *Aethionema grandiflorum, Alliaria petiolata, Cardamine matthioli, Chamira circaeoides*, and *Rorippa amphibia* were downloaded from Genbank.

### Analyses

#### Drought frequency calculations

To characterize drought regimens across the distributions of annual and perennial species of *Heliophila*, we calculated drought during different seasons at the location of observations for *Heliophila* records using the VHI. Specifically, we created global maps of the frequencies of observing drought conditions (VHI<40, NOAA) during the winter (quarter surrounding winter solstice), spring (quarter surrounding spring equinox), summer (quarter surrounding summer solstice) and fall (quarter surrounding fall equinox) from 1981 to 2015. From these maps, the drought frequency during the winter, spring, summer, and fall were extracted for the locations of all GBIF records.

#### Filtering of occurrence records

To avoid instances with spurious location data, we filtered raw GBIF by restricting our analyses to include only:

- records for species with reported life history
- records with geospatial data
- records without known geospatial coordinate issues (i.e., coordinates reported are those of herbarium)
- records from collection sites classified as land pixels in the VHI dataset
- records from Africa (to exclude locations of cultivation)
- records without duplicates (i.e., identical species, location, collection date)

#### Phylogeny construction

Out group (*Aethionema grandiflorum, Alliaria petiolata, Cardamine matthioli, Chamira circaeoides*, and *Rorippa amphibia*) and ingroup *Heliophila* ITS I and II sequences were aaligned using MAFFT (Katoh *et al*., 2002) with strategy G-INS-I, offset value 0.1, and all other options set as default. The *GTR* + Γ model of nucleotide substitution was determined to best fit the data based on AIC using jModelTest2 (Guindon & Gascuel, 2003; Darriba *et al*., 2012). A maximum clade credibility tree with branch lengths as relative time was estimated by summarizing data from six runs of 100,000,000 generations of Bayesian Markov chain Monte Carlo conducted in BEAST 2 (Bouckaert *et al*., 2014). Model selection and phylogenetic analyses were conducted through the CIPRES Science Gateway (Miller *et al*., 2010).

#### Comparison of drought frequency between annual and perennial species

To evaluate the hypothesis that annual and perennial life history strategies reflect adaptations to alternative drought regimes, we tested the corresponding prediction that the observed distributions of annual and perennial *Heliophila* species would be significantly associated with historic drought frequency. First, we compared the frequency of drought during the winter, spring, summer, and fall between total occurrence records of annual and perennial species by t-tests. To account for variation in the number of occurrence records per species, we next calculated the mean drought frequency during the winter, spring, summer and fall for each species. Because shared evolutionary history of closely related species can lead to spurious associations between traits and environments (Felsenstein, 1985), we tested for a relationship between life history strategy and drought frequency while controlling for phylogeny using phylogenetic logistic regression (Ives & Garland, 2010).

#### Collection dates

To test the hypothesis that annual species have adapted to escape drought prone seasons as seeds, collection dates for herbarium specimens were compared between annual and perennial species. Comparisons of distributions were made by Two-sample Kolmogorov-Smirnov test and Barlett variance test.

## Results

Out of 8670 *Heliophila* GBIF records, 6634 were for species with reported life history (Mummenhoff *et al*., 2005), 2856 had geospatial data, 2833 did not have geospatial issues, 2684 were located on pixels classified as land having drought measurements, 2543 were located in Africa, 2192 were not duplicated. Thus, after all filtering steps, 2192 records for 42 species (Figure 1, Table S1) passed for further analyses. The number of samples varied between species, with a mean of 52.19 samples per species. *H. rigidiuscula* had the most records, 201, and *H. cornellsbergia* the fewest, 2 (Table S1).

**Figure 1.**
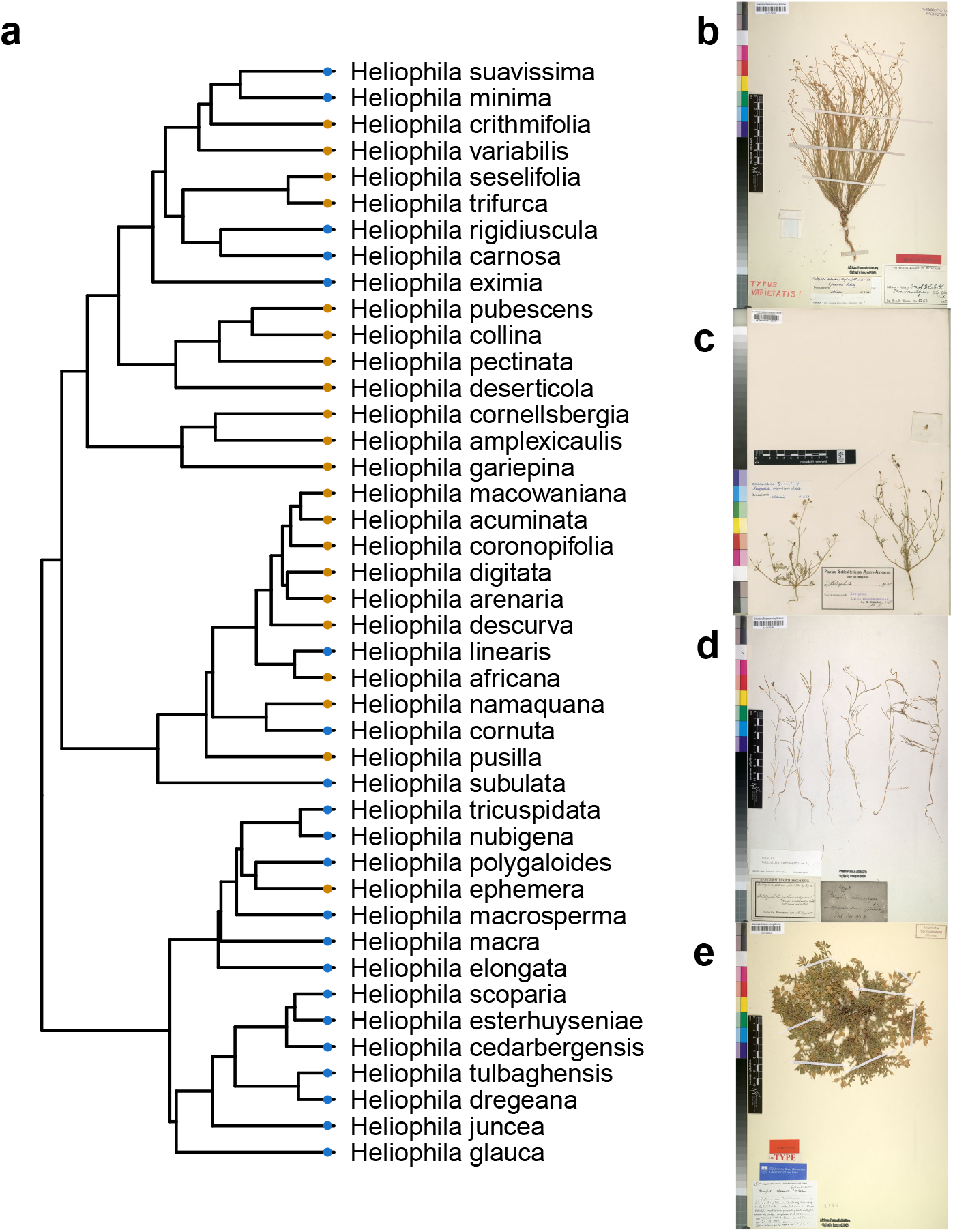
Species and examples of herbaria specimens of *Heliophila* (a) Phylogeny and life history strategies of species studied. Orange circles at branch tips mark annual species and blue circles mark perennial species. Example herbaria specimens accessed via GBIF of (a) *H. minima*, (b) *H. deserticola*, (c) *H. coronopifolia* and (d) *H. ephemera*. Images (a,c,d) courtesy of The Bavarian Natural History Collections (CC BY-SA 4.0) and (b) The London Natural History Museum (CC BY 4.0). Links to images are found in the supplement.

There were clear visual differences between the distributions of the 960 annual and the 1232 perennial *Heliophila* observation records (see Figure S1 for maps of individual species). While annual species were generally found in the western regions of South Africa and Namibia, primarily in the Cape Floristic Region and Succulent Karoo (Figure 2a), the occurrence of perennials extended to the east coast of South Africa (Figure 2b).

**Figure 2.**
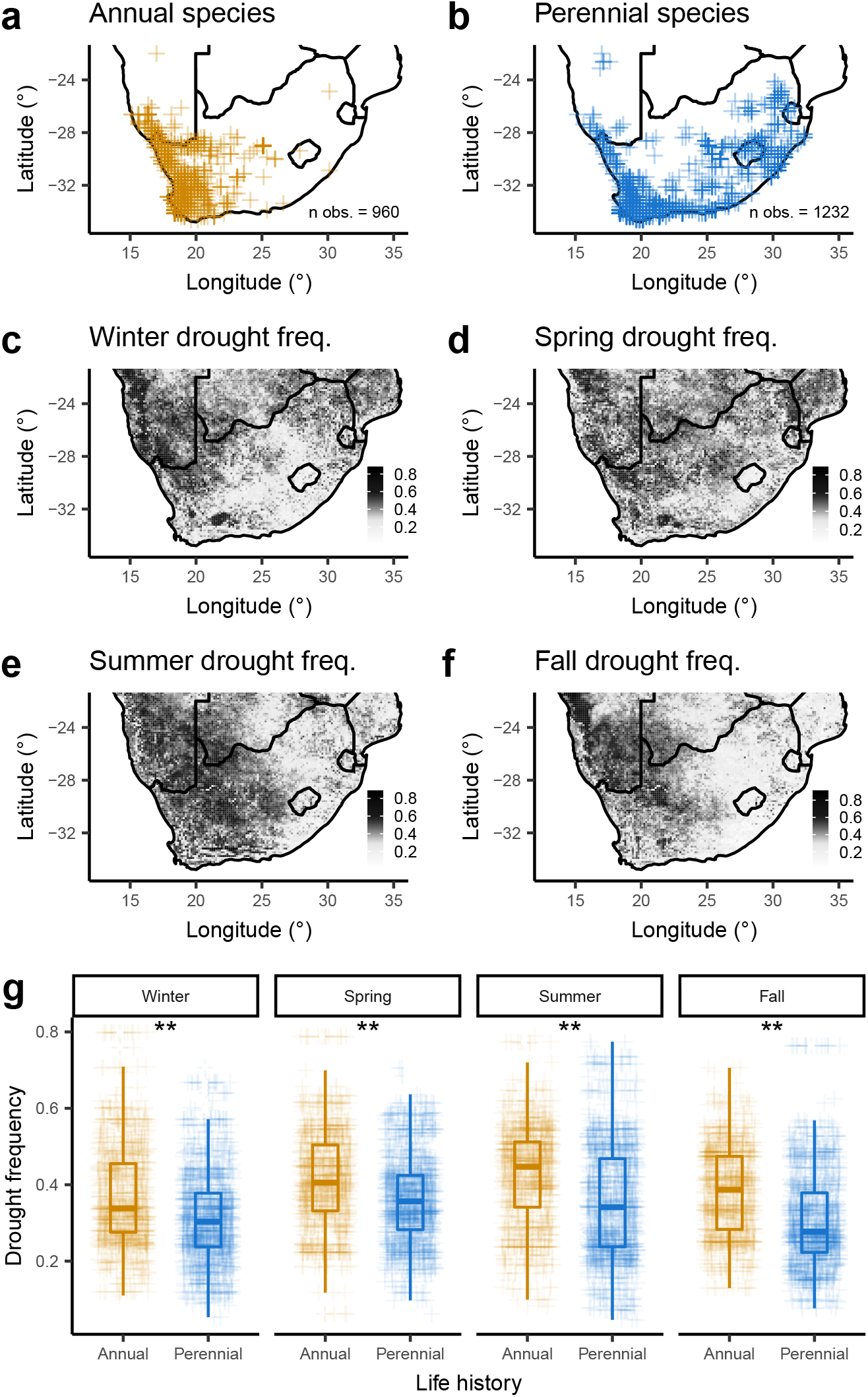
Locations of occurrence records of (a) annual and (b) perennial *Heliophila*. Drought frequency during the (c) winter, (d) spring, (e) summer and (f) fall measured using the VHI. (g) Drought frequencies during each season at the observation locations of annual and perennial *Heliophila* (t tests, ** = p < 0.01).

The frequency of drought varied considerably across the ranges of *Heliophila* species (Figure 2c-f). This heterogeneity is expected, given that this is one of the most climatically diverse regions of the Earth (Sayre *et al*., 2013). It is worth noting the east to west cline in drought frequency observed during the summer, which distinguishes the high drought frequency of the Kalahari Sands and Namid Desert phytogeographic regions from the low drought frequency of the Drakensberg Mountains and Coastal Zambesian phytogeographic regions. In the Cape phytogeographic region there was finer scale heterogeneity in drought frequency during the summer.

Theory predicts that annuality should be adaptive in places where stresses such as drought are more common. Conversely, perenniality should be adaptive in places where such stresses are less frequent. We found that the frequency of drought was significantly higher at the locations of occurrence records for annual species. When comparing across all occurrence records (all records rather than species means, Figure 2g), the frequency of drought was significantly higher at the location of annuals during the winter (t = 10.65, p = 0.00), spring (t = 10.73, p = 0.00), summer (t = 12.67, p = 0.00), and fall (t = 15.26, p = 0.00). Because a comparison across all occurrence records does not account for variation in the number of records per species (Table S1) or species relatedness (Figure 1a), we also tested whether mean drought frequency values of each species were significantly different between annuals and perennials using phylogenetic logistic regression. We found that the mean drought frequencies were significantly higher (*α* = 0.05) in annual species during the spring, summer, and fall (Table 1, Figure 3a). These findings indicate that common acestry alone does not explain differences the drought frequencies experienced between the environments of annual and perennial *Heliophila*.

**Figure 3.**
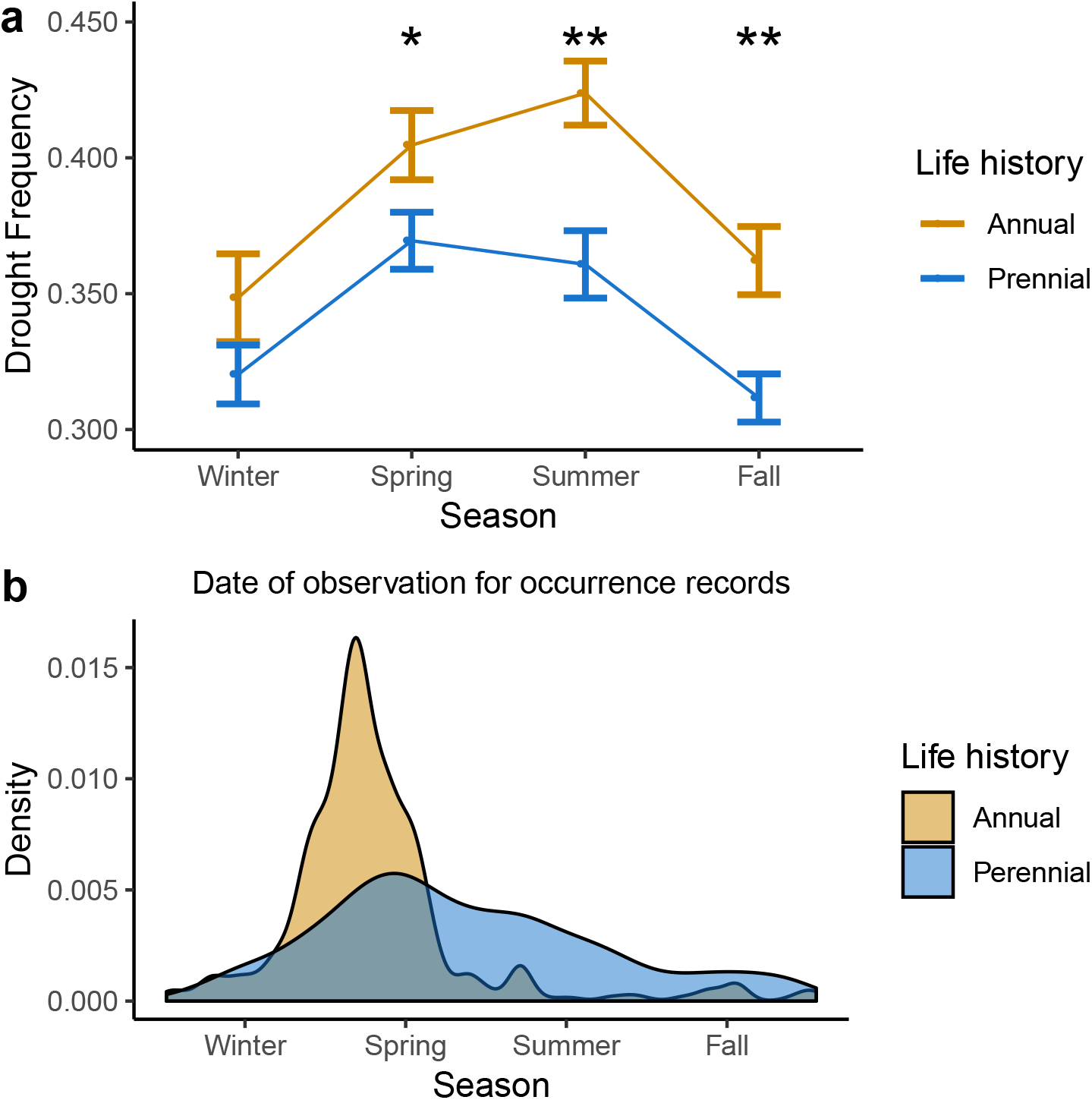
(a) Comparison (mean +− SE) of drought frequency across seasons measured at the GBIF records of annual and perennial species of *Heliophila*. (phylogenetic logistic regression, * = p < 0.05, ** = p < 0.01) (b) Collection dates of GBIF records of annual and perennial species of *Heliophila*.

**Table 1.**
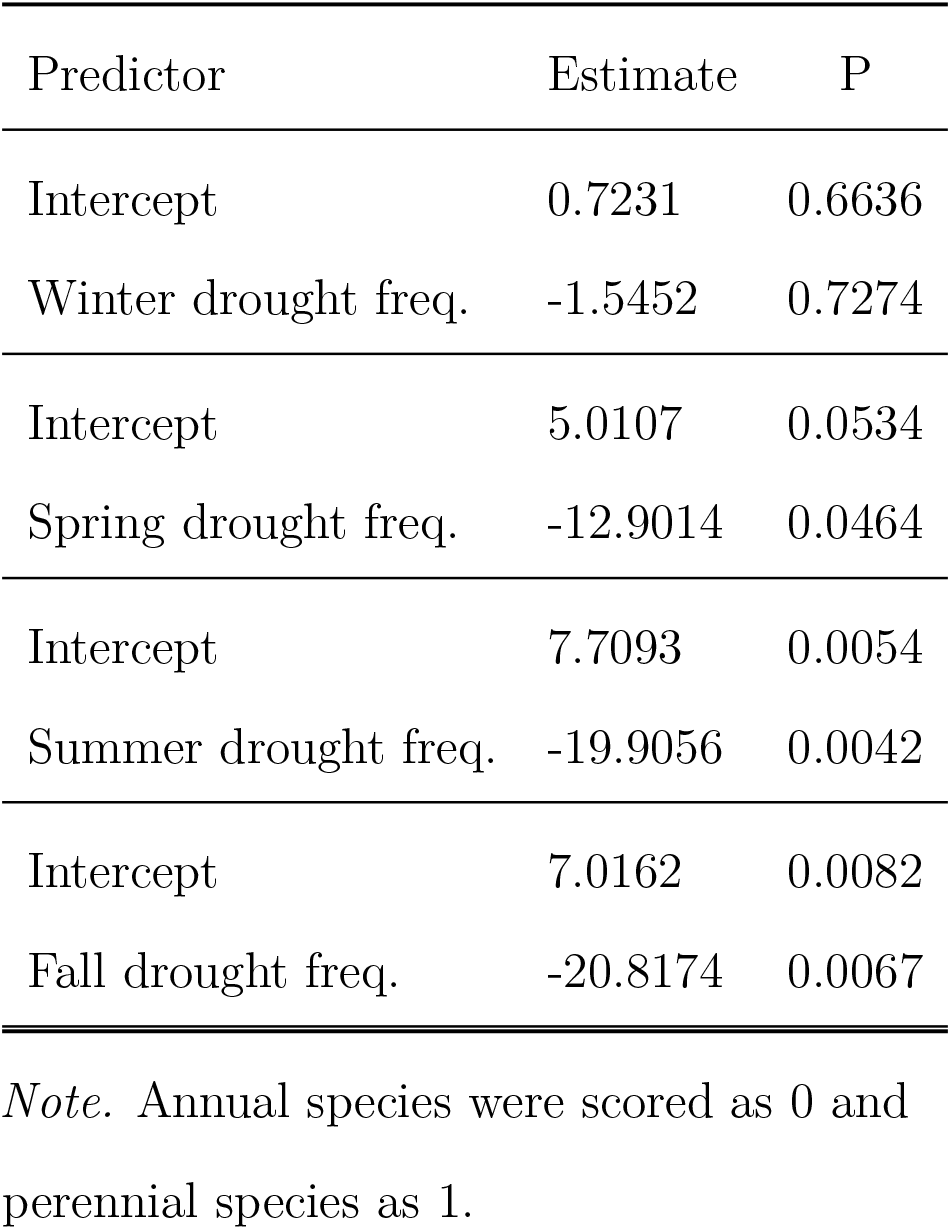
*Phylogenetic logistic regressions between life history, and the mean drought frequency observed at specimen sites of Heliophila species the winter, spring, summer, and fall*.

| Predictor | Estimate | P |
| --- | --- | --- |
| Intercept | 0.7231 | 0.6636 |
| Winter drought freq. | -1.5452 | 0.7274 |
| Intercept | 5.0107 | 0.0534 |
| Spring drought freq. | -12.9014 | 0.0464 |
| Intercept | 7.7093 | 0.0054 |
| Summer drought freq. | -19.9056 | 0.0042 |
| Intercept | 7.0162 | 0.0082 |
| Fall drought freq. | -20.8174 | 0.0067 |

*Note.* Annual species were scored as 0 and perennial species as 1.

The preceding results indicate that annual species are found in environments where droughts are significantly more frequent, especially in the summer and fall. Classic life history theory hypothesizes that annuality reflects adaptation to such environments because it allows species to escape stressful conditions. If this is the case, we would expect that annuals spend the drought prone seasons of summer and fall as seeds. To test this hypothesis, we compared the dates of occurrence records between annual and perennial *Heliophila* species. The distributions reveal a considerable difference in the timing of observation of these two life histories. In comparison to perennials, which appear to be collected throughout the year, annuals are almost exclusively observed during the winter and spring (Figure 3b). The differences between the distribution of collection dates were significant by all tests (ks.test D = 0.25, p = 0; bartlett.test K2 = 503.18, p = 0.00) This is consistent with a model of life history in which annual species flower in the spring, set seed, senesce, and die before the summer. Thus, these annual species are likely to remain dormant during the summer and fall, when drought is the strongest predictor of the distributions of annual and perennial life histories (Figure 3a).

## Discussion

To test the hypothesis that annual and perennial plants reflect adaptation to alternative drought environments we examined the landscape distribution of life history strategies in the large and diverse mustard genus, *Heliophila*. Using metadata of 2192 occurrence records and a 34 year dataset of satellite-detected droughts, we tested the prediction that annual species are more often observed in drought-prone locations than perennial species, when controlling for phylogenetic relatedness. We found that drought frequency is significantly different between the distributions of annual and perennial species, with annuals being found in environments with more frequent drought, and that this signal is strongest during the seasons when annuals are likely escaping via seed dormancy. These results remain significant while controlling for the phylogenetic relationships of *Heliophila* species, yielding support for the role that natural selection has played in driving contemporary distributions of these alternatives strategies in relation to drought regimens.

We cannot eliminate the possibility that confounding traits or environmental variables are the causative factors explaining variation in the distributions of annual and perennial species. Nevertheless, these results provide quantitative support for the classic prediction that annual species are found in environemnts that experience more frequent drought than perennial species. These findings complement previous reports of qualitative associations between annuality with enviroments characterized as having increased aridity (Evans *et al*., 2005), alternative precipitation defined habitats (Morishima *et al*., 1984; Datson *et al*., 2008), or greater unpredictability (Cruz-Mazo *et al*., 2009). However, to our knowledge this is the first study to demonstrate a significant association between life history and drought in a phylogenetic context informed by large scale species distribution data and long term drought measures.

Unfortunately, herbarium collections and their associated data do not represent systematic or random sampling of a species distribution. Significant biases in collecting exist, which we have not necessarily controlled for here, and may have some effect on our findings, such as a bias toward collecting near roads or near the locations of natural history collections (Daru *et al*., 2018). Future research will benefit from systematic sampling efforts to avoid these noted biases. However, the ecosystems of southern Africa include several biodiversity hotspots and are among the most botanically well sampled regions on Earth (Daru *et al*., 2018), suggesting that this may currently be the optimal region for our analyses of life history distribution. Indeed, we were able to use 2192 occurrence records to study 42 species, which represents a significant advance over relying on personal observations to characterize species distributions.

These findings support classical theoretical predictions about the adaptive value of annual and perennial life history strategies. Taken together, they suggest that in *Heliophila*, annual species are adapted to environments with increased summer droughts by avoiding these seasons in a dormant seed phase of their life cycle. They also suggest that perenniality is adaptive in environemnts where droughts are less frequent. While most previous work has focused on describing the evolutionary origins of annuality (Barrett *et al*., 1996; Conti *et al*., 1999; Andreasen & Baldwin, 2001; Verboom *et al*., 2004; Friedman & Rubin, 2015) there are at least a few other cases where perenniality appears to have arisen from an annual ancestor (Bena *et al*., 1998; Tank & Olmstead, 2008). And while early theory predicted selection for annuality when adult morality is high (Stearns, 1992), we also find evidence that perenniality could be explained by reduced frequency of drought. The phylogeny reveals several transitions from annual to perennial life history (Figure 1a) and the distributions of perennial *Heliophila* extend into regions where drought frequency is low (Figure 2b, Figure S1). Perennials may be able to out complete annual relatives in environments where the infrequency of drought favors strategies that allow plants to benefit from growth over many seasons. This also suggests that annuals rely on drought as a source of disturbance for seedling recruitment when competing with perennials (Corbin & D’Antonio, 2004). Indeed, no annual species were observed in the low drought regions of eastern South Africa (Figure 2, Figure S1).

These findings suggest that species with locally adaptive life history strategies could be threatened by rapidly changing drought regimens (Dai, 2011). This could have impacts on ecosystem functioning and processes such as carbon cycling if life history traits evolve or the composition of annual and perennial species changes in response (Garnier *et al*., 1997; Roumet *et al*., 2006; Monroe *et al*., 2018a). Furthermore, the frequency of drought may be an important factor when considering the use of perennial cropping systems (Parry *et al*., 2005; Lelièvre & Volaire, 2009).

In conclusion, we find strong support for classic life history theory which predicts that annuality is adaptive in environments where droughts occur more frequently. Additionally, we report evidence consistent with a life history model in annuals in which they escape drought prone seasons during the seed phase of their life cycle. Finally, we find evidence that the distributions of perennial lineages may indicate a competitive advantage in areas where droughts are infrequent. More broadly, this work highlights the irreplaceable value of natural history collections and demonstrates the power of combining such information with large scale remote sensing data to address outstanding classic hypotheses in ecology and evolution.

## Acknowledgments

We thank members of the Sloan lab for generous feedback that improved the quality of this work. This research was supported by NSF Award 1701918 and USDA-NIFA Award 2014-38420-21801 to JGM.

## Author contributions

JGM, BG, KGT and JKM contributed to the design of the research, interpretation, and writing the manuscript. JGM, BG, and KGT contributed to the performance of the research and data analysis.

## Supplement

### Images used

https://www.gbif.org/occurrence/1099023487 https://www.gbif.org/occurrence/1057389408 https://www.gbif.org/occurrence/1099023562 https://www.gbif.org/occurrence/1099023490

### Software used

We used R (Version 3.5.1; R Core Team, 2018) and the R-packages *ape* (Version 5.2; Paradis & Schliep, 2018; Orme *et al*., 2018; Soetaert, 2018), *bindrcpp* (Version 0.2.2; Müller, 2018), *caper* (Version 1.0.1; Orme *et al*., 2018), *coda* (Version 0.19.2; Plummer *et al*., 2006), *diagram* (Version 1.6.4; Soetaert, 2017), *dplyr* (Version 0.7.8; Wickham *et al*., 2018), *forcats* (Version 0.3.0; Wickham, 2018a), *gee* (Version 4.13.19; R by Thomas Lumley & author., 2015), *geiger* (Version 2.0.6; Alfaro *et al*., 2009; Harmon *et al*., 2008; Eastman *et al*., 2011; Slater *et al*., 2012), *ggplot2* (Version 3.1.0; Wickham, 2016), *logistf* (Version 1.23; Heinze & Ploner, 2018), *maps* (Version 3.3.0; Richard A. Becker *et al*., 2018), *MASS* (Version 7.3.51.1; Venables & Ripley, 2002), *Matrix* (Version 1.2.15; Bates & Maechler, 2018), *MCMCglmm* (Version 2.26; Hadfield, 2010), *mvtnorm* (Version 1.0.8; Genz & Bretz, 2009), *papaja* (Version 0.1.0.9842; Aust & Barth, 2018), *phylolm* (Version 2.6; Ho & Ane, 2014), *phytools* (Version 0.6.60; Revell, 2012), *purrr* (Version 0.2.5; Henry & Wickham, 2018), *raster* (Version 2.8.4; Hijmans, 2018), *readr* (Version 1.2.1; Wickham *et al*., 2017), *shape* (Version 1.4.4; Soetaert, 2018), *sp* (Version 1.3.1; Pebesma & Bivand, 2005), *stringr* (Version 1.3.1; Wickham, 2018b), *tibble* (Version 1.4.2; Müller & Wickham, 2018), *tidyr* (Version 0.8.2; Wickham & Henry, 2018), and *tidyverse* (Version 1.2.1; Wickham, 2017) for all our analyses.

## Supplementary tables and figures

**Table S1.**
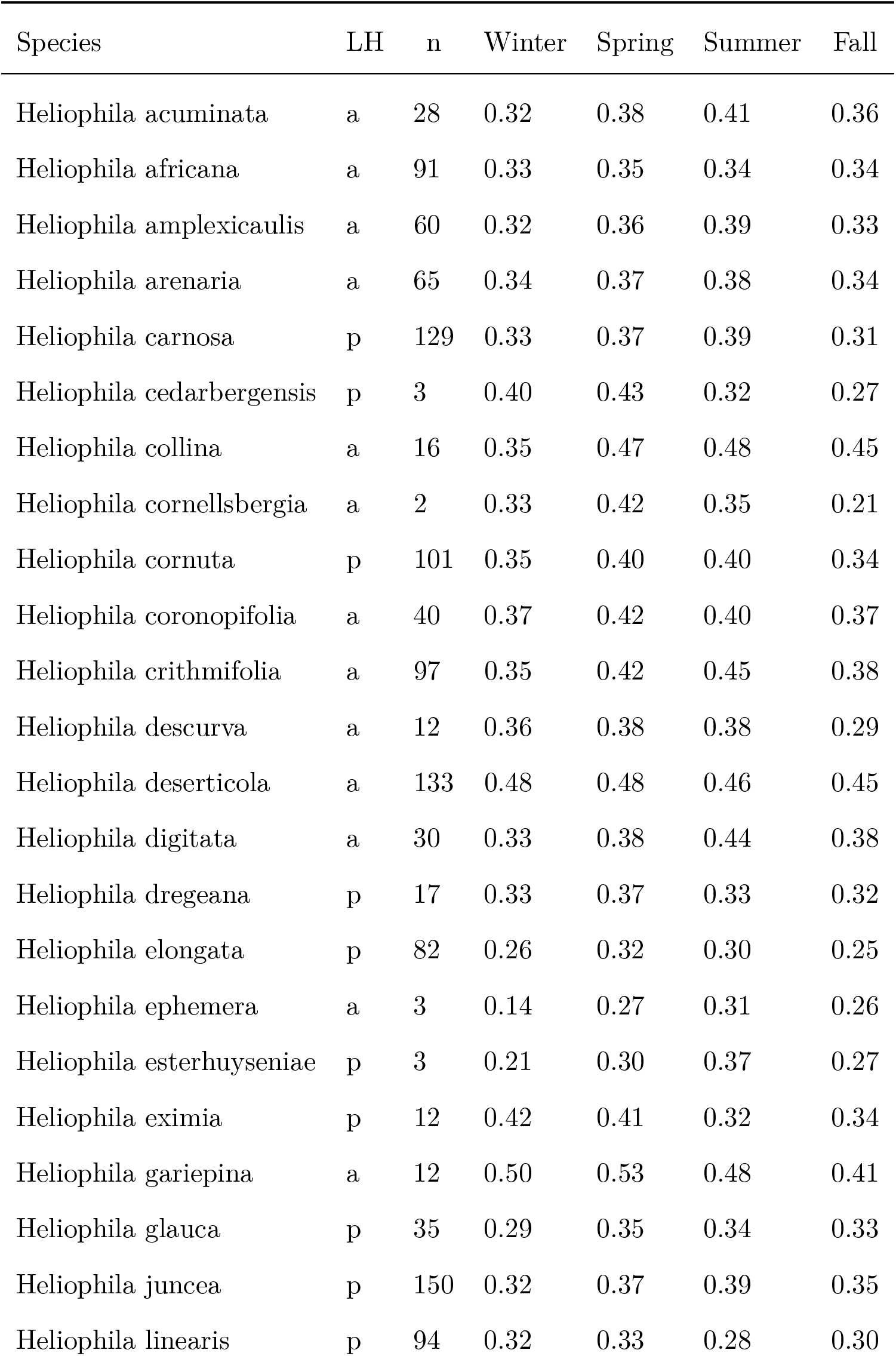
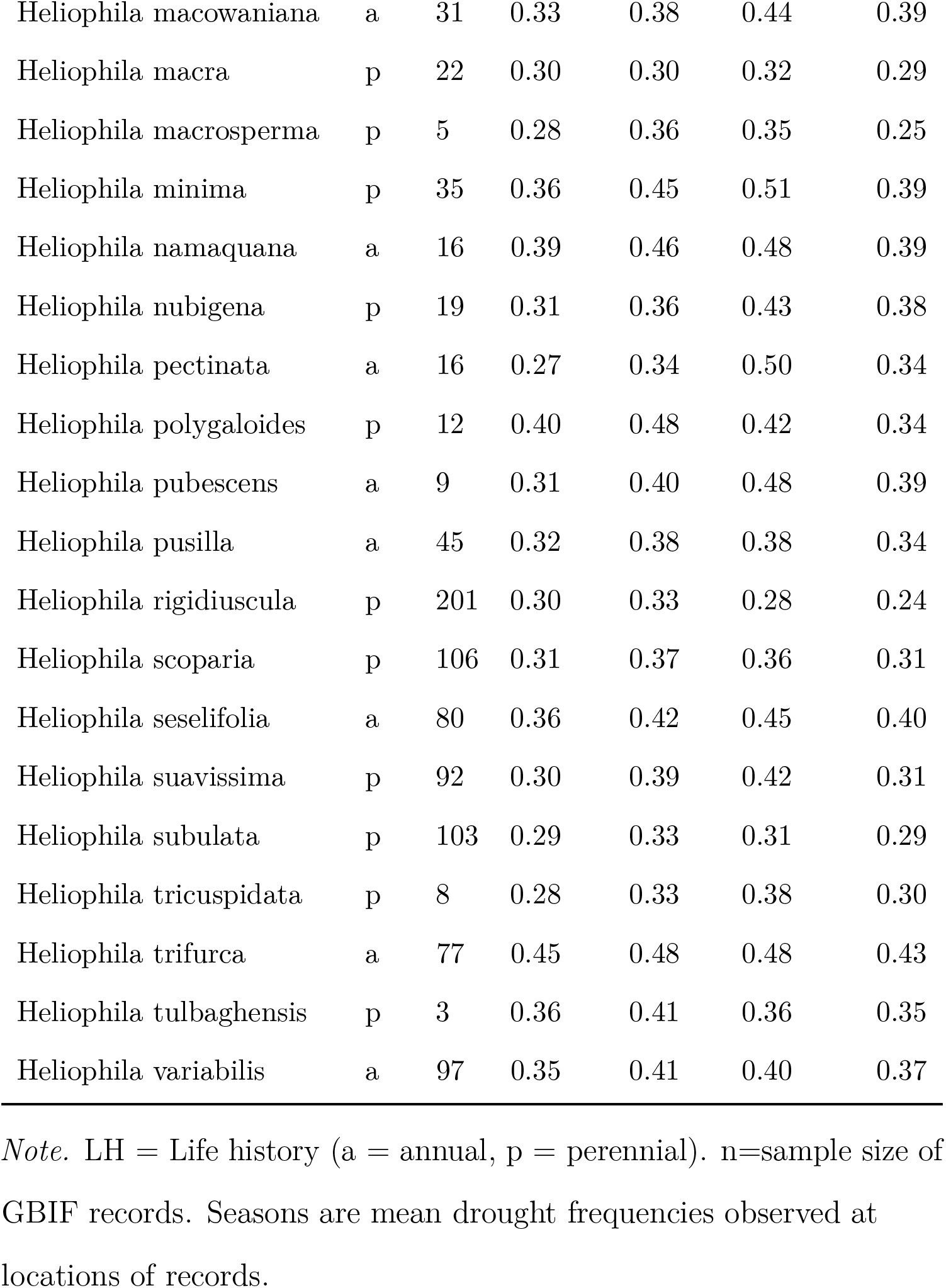
*Heliophila species records and the mean drought frequencies during different seasons at the location of records*

**Figure S1.**
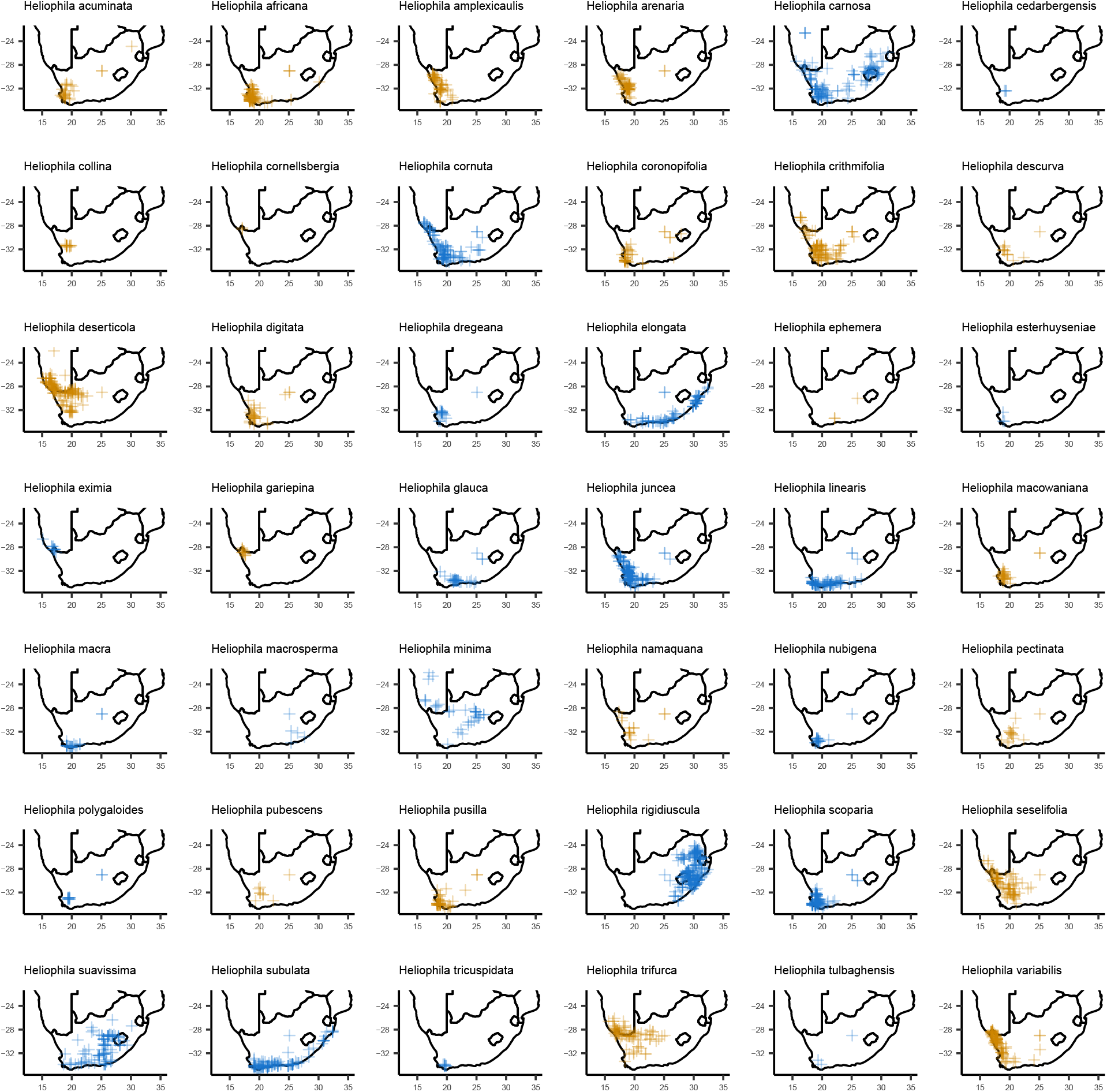
Maps of occurrence records for individual species. Orange points indicate annual species. Blue points indicate perennial species.

